# DATRASextra: An R package for streamlined workflows with ICES DATRAS bottom-trawl survey data

**DOI:** 10.64898/2026.06.29.735240

**Authors:** Tobias K. Mildenberger, Federico Maioli, Casper W. Berg

## Abstract

Scientific bottom-trawl surveys provide essential fisheries-independent data for fisheries and ecosystem research. In the Northeast Atlantic, the ICES Database of Trawl Surveys (DATRAS) compiles haul-level information, species- and length-specific catch data, and individual biological observations across multiple long-term surveys. However, reproducible workflows for processing and integrating these relational datasets remain challenging. We present `DATRASextra`, an open-source R package that provides modular end-to-end workflows for accessing, cleaning, harmonising, quality-controlling, and analysing DATRAS survey data. The package supports derivation of standardised haul-level survey variables, integration of multiple surveys, and generation of analysis-ready datasets for downstream applications including stock assessment, biodiversity analyses, and large-scale synthesis efforts such as Fish-Glob.

## 1. Motivation and significance

Fisheries-independent data from scientific marine bottom-trawl surveys are central to monitoring fish communities across space and time. These surveys provide standardised information on species abundance, distribution, population structure, and biological characteristics, supporting fisheries stock assessments, biodiversity monitoring, and ecosystem analyses. In the North-east Atlantic and adjacent seas, the International Council for the Exploration of the Sea (ICES) Database of Trawl Surveys (DATRAS) serves as a central repository for such data, currently compiling information from 28 surveys spanning more than 60 years [1].

DATRAS data underpin a broad range of scientific and advisory applications. In fisheries science, survey indices derived from DATRAS are routinely used to estimate trends in stock abundance, recruitment, and population structure for most of the 271 stocks. Beyond single-stock assessment, the database also supports biodiversity assessments [e.g. 2], ecosystem-based fisheries management [e.g. 3, 4], and large-scale ecological syntheses [e.g. 5, 6, 7]. Recent examples include the FishGlob initiative, which integrates bottom-trawl survey data across regions to investigate global patterns in marine fish communities [8, 9].

**Table 1:** Code metadata (mandatory)

| Nr. | Code metadata description | Metadata |
| --- | --- | --- |
| C1 | Current code version | v0.4.0 |
| C2 | Permanent link to code/repository used for this code version | https://github.com/tokami/DATRASextra/releases/tag/v0.4.0 |
| C3 | Legal Code License | GNU General Public License (GPL) v3 |
| C4 | Code versioning system used | Git |
| C5 | Software code languages, tools, and services used | R |
| C6 | Compilation requirements, operating environments & dependencies | R ($\geq 4.0.0$), https://github.com/DTUAqua/DATRAS |
| C7 | If available Link to developer documentation/manual | https://tokami.github.io/DATRASextra |
| C8 | Support email for questions |  |

Despite standardised data formats and internal validation procedures within DATRAS, practical workflows for accessing and processing these data remain complex. Users typically need to download and merge multiple relational tables containing haul-level, length-based, and biological observations; apply survey-specific filtering and harmonisation procedures; implement quality-control steps; and construct reproducible analysis-ready datasets. These tasks can require extensive project-specific scripting and substantial familiarity with both the DATRAS database structure and survey-specific conventions. As a consequence, workflows are often difficult to reproduce, compare, and maintain across studies and institutions. These challenges have motivated recent efforts to establish best-practice guidelines for survey data processing and integration [e.g. 10].

The scale and heterogeneity of the DATRAS database further increase the complexity of reproducible analyses. The database currently contains information from approximately 142,000 hauls, around 2,000 Aphia IDs, and more than 20 million biological records distributed across multiple linked tables. Some survey time series extend back to 1965, resulting in substantial variation in sampling protocols, spatial coverage, taxonomic resolution, and survey design over time.

Existing R packages such as `icesDatras` [11] and `DATRAS` [12] provide important infrastructure for accessing and handling DATRAS data. However, these packages primarily focus on data retrieval and basic manipulation and do not provide integrated end-to-end workflows for data cleaning, quality control, harmonisation across surveys, the generation of standardised analysis-ready products, or comprehensive documentation. In other regions, comparable efforts such as the surveyjoin package [13] provide standardised access to multiple scientific trawl surveys in the Northeast Pacific, but no equivalent integrated, end-to-end toolbox currently exists for the ICES DATRAS data. `DATRASextra` was developed to address these limitations by providing a user-friendly and well-documented toolbox for reproducible DATRAS end-to-end workflows in R. The package supports the complete workflow from raw data access to analysis-ready outputs, including harmonisation of surveys and species information, reproducible quality-control procedures, calculation of stratified mean abundance and biomass indices, and generation of standardised survey products. In addition, the package includes vignettes demonstrating common workflows and reproducible examples, including the reproduction of the Northeast Atlantic component of the FishGlob dataset [8], comprising approximately 45% of the included surveys.

Typical workflows in `DATRASextra` begin with downloading raw survey data directly from DATRAS, followed by automated merging of relational tables, implementation of quality-control procedures, and generation of standardised survey products such as abundance or biomass indices. The resulting datasets can then be directly used in downstream applications including stock assessments, biodiversity analyses, and species distribution modelling. By consolidating these steps into a single reproducible framework, `DATRASextra` reduces the complexity of working with DATRAS data and facilitates transparent and standardised analyses across users and applications.

## 2. Software description

### 2.1. Software architecture

`DATRASextra` is organised as a modular workflow-oriented framework for accessing, processing, quality-controlling, and analysing ICES DATRAS survey data in R [14]. The package builds on the ICES DATRAS web services and existing infrastructure provided by the icesDatras and DATRAS packages, while extending these with integrated end-to-end processing workflows.

The overall software architecture is illustrated in Figure 1. The workflow is structured into four main components: (i) data ingestion, (ii) core processing and harmonisation, (iii) quality control and enrichment, and (iv) generation of analysis-ready outputs.

**Figure 1:**
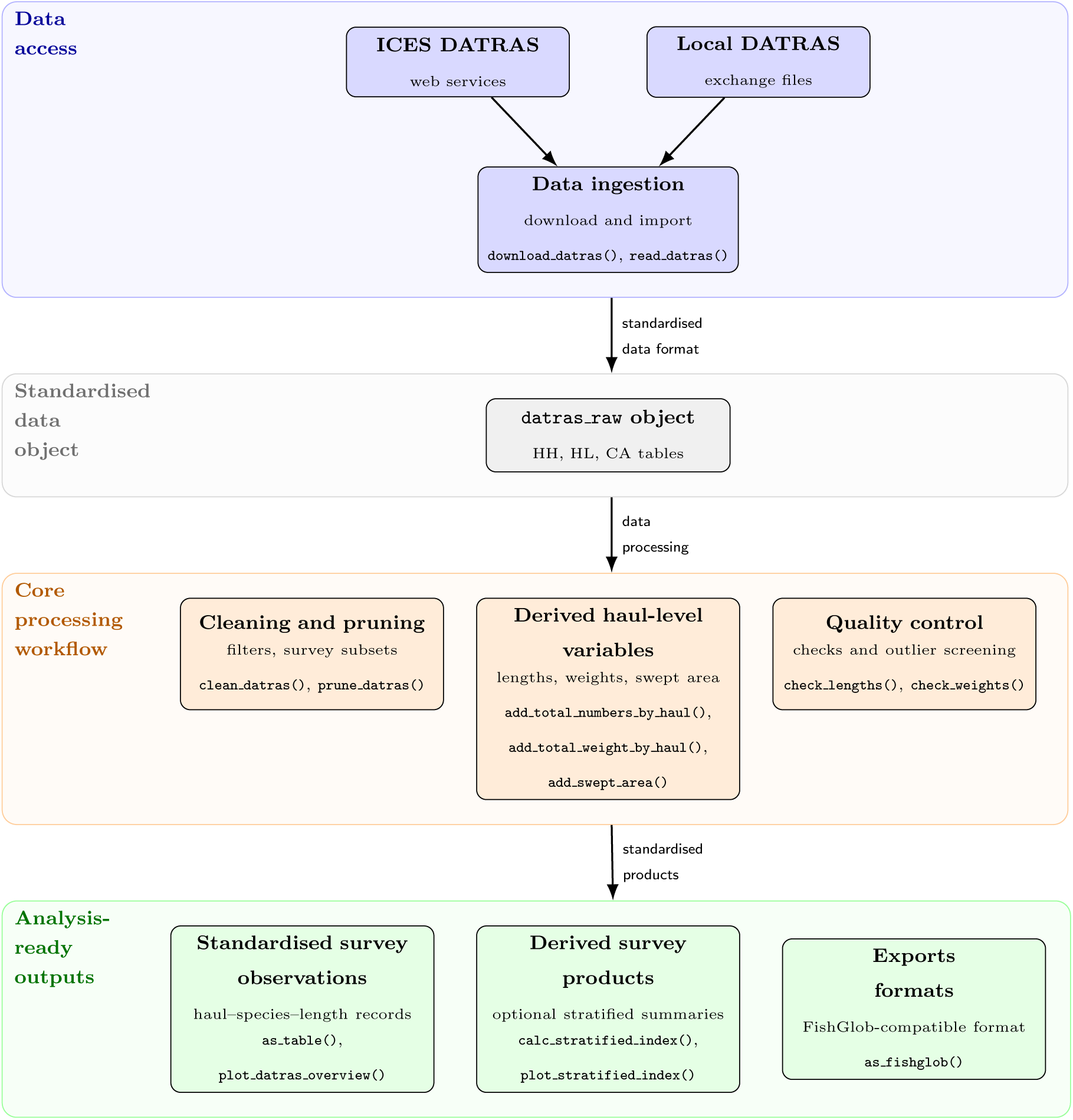
Overview of the modular workflow architecture implemented in `DATRASextra`. Data are imported from ICES DATRAS web services or local exchange files, organised into a standardised datras raw object, processed through modular cleaning, enrichment, and quality-control routines, and exported as analysis-ready products for downstream ecological and fisheries applications.

Data can be imported either directly from the ICES DATRAS API or from locally archived exchange files. Downloaded data are organised into a standardised `datras_raw` object containing haul-level (HH), length-based (HL), and biological (CA) data tables. Core processing functions then support harmonisation across surveys, species, years, and quarters, as well as filtering, pruning, and integration of auxiliary information such as swept-area estimates and weight-at-length relationships.

The package further includes modular quality-control routines for identifying invalid hauls, inconsistent biological measurements, and statistical outliers. Processed datasets can subsequently be exported into multiple standardised output formats suitable for downstream analyses, including abundance and biomass indices, visual summaries, and FishGlob-compatible products.

A central design principle of `DATRASextra` is reproducibility through explicit workflow functions and standardised intermediate objects. The modular architecture allows users to combine individual components flexibly while maintaining transparent and reproducible analysis pipelines. Internal reference datasets, including species and survey metadata, are distributed with the package to ensure consistency across workflows.

### 2.2. Software functionalities

The major functionalities of `DATRASextra` span the full survey workflow and operate on the standardised `datras_raw` object introduced above, which is processed through a series of interoperable workflow functions.

Core functionality includes automated downloading and local archiving of complete or user-defined subsets of DATRAS data through the ICES DATRAS web services. The package further provides functions for importing and harmonising exchange files and relational survey tables into reproducible analysis workflows.

A central functionality of `DATRASextra` is the integration of information distributed across multiple DATRAS tables to derive standardised haul-level survey variables. This includes the calculation of swept area from haul metadata, derivation of numbers-at-length and biomass-at-length observations, aggregation of catch information by custom length classes, and conversion of individual biological observations into analysis-ready haul-level summaries. These workflows allow users to transform relational survey data into coherent standardised survey observations suitable for downstream analyses.

The package also includes functions for data cleaning, harmonisation, and quality control. These include survey and species filtering, taxonomic standardisation, outlier screening, consistency checks for biological measurements, and haul-level diagnostics. Quality-control workflows are designed to support transparent and reproducible preprocessing of large heterogeneous survey datasets.

In addition, `DATRASextra` provides tools for integrating multiple surveys and producing standardised export formats suitable for downstream ecological and fisheries applications, including FishGlob-style workflows [8]. Flexible plotting and exploratory visualisation functions support rapid inspection of survey coverage, haul distributions, and processed survey products.

The package is accompanied by extensive documentation, including function-level help pages, reproducible tutorials, and workflow-oriented vignettes. These demonstrate typical use cases ranging from downloading and preprocessing raw DATRAS data to generating standardised survey datasets for stock assessment, biodiversity analyses, and species distribution modelling. Full documentation is available through the package pkgdown website and included vignettes.

## 3. Illustrative examples

The following examples demonstrate typical workflows implemented in `DATRASextra`, including a minimal end-to-end workflow, derivation of length-structured haul-level survey variables, and reproduction of parts of the FishGlob data-processing pipeline. The examples require the package to be installed, for example with:

~~~
remotes::install_github(“tokami/DATRASextra”)
~~~

and loaded into the R environment with:

~~~
library(DATRASextra)
~~~

### 3.1. Example 1: Minimal workflow

The following example demonstrates how to download, process and visualise DATRAS survey data using a concise reproducible workflow. Specifically, it illustrates spatial and temporal variation in haul-level catch rates of Atlantic herring (*Clupea harengus*) from the North Sea International Bottom Trawl Survey (NS-IBTS).

~~~
## Download NS-IBTS survey data (1991, 2001, 2011, 2021)s
dat <-download_datras(surveys = “NS-IBTS”, years = c(1991, 2001, 2011, 2021))
## Apply standard processing and subset Atlantic herring her <-clean_datras(dat, aphias = “126417”)
## Derive total abundance per haul her <-add_total_numbers_by_haul(her)
## Plot haul-level catches in space and time plot_datras_overview(her, metric = “mean”, value_var = “HaulN”, transform = “sqrt”, by_year = TRUE, multi_panels = TRUE)
~~~

The results show a patchy distribution of hauls with high catch rates and an aggregation of high catch rates in waters between Denmark and Sweden in more recent years (Figure 2). These haul-level quantities, including total abundance and standardised catch rates, can be used directly as input to downstream analyses such as abundance-index standardisation, stock assessment, biodiversity analyses, or species distribution modelling. With minor modifications, the same workflow can be extended to specific length or age groups, biomass-based quantities, multiple species, or combined survey datasets. Additional use cases and extended workflows are provided in the package vignettes.

**Figure 2:**
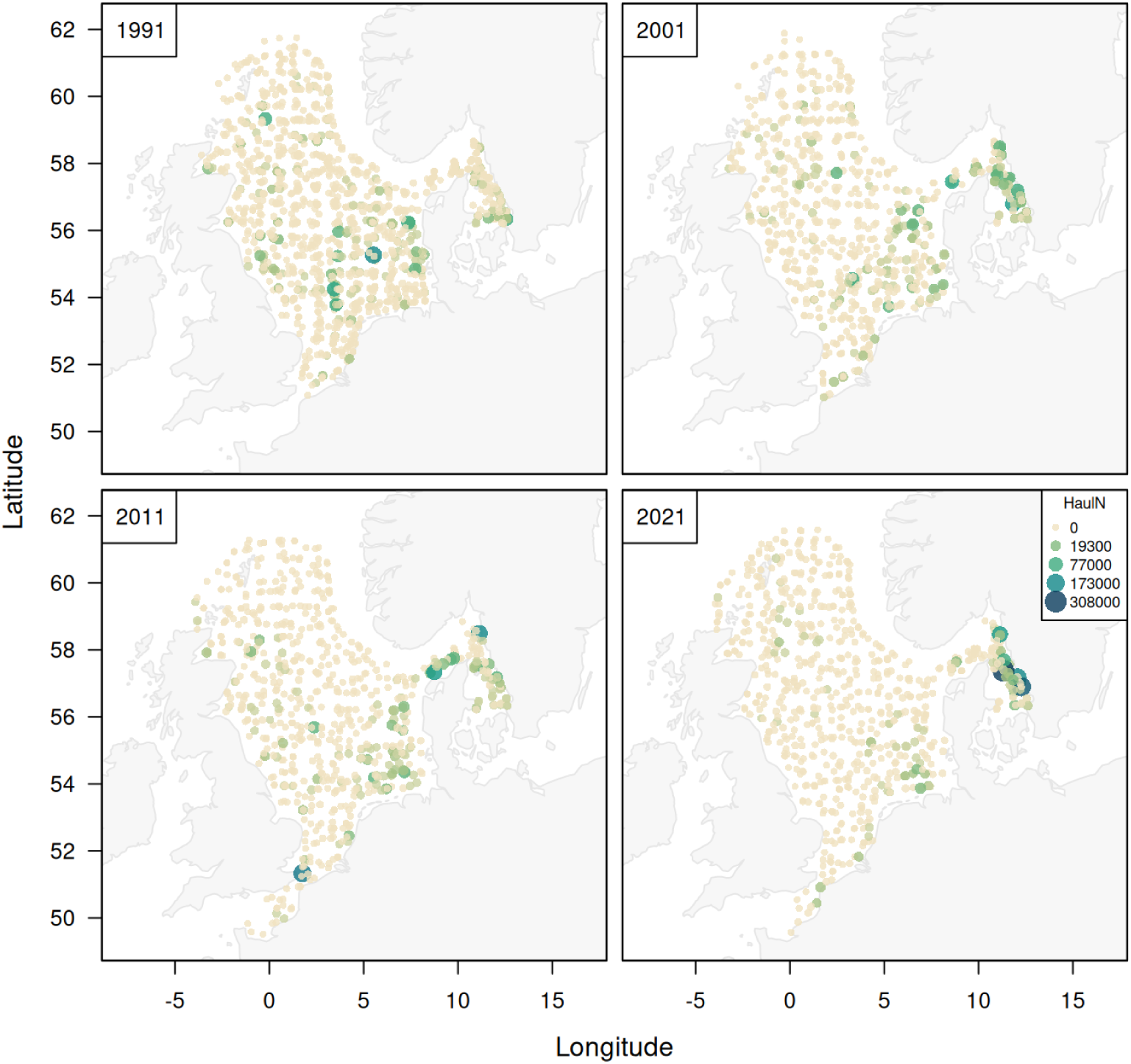
Example output generated with `DATRASextra` showing the spatial distribution of Atlantic herring (*Clupea harengus*) catch rates from the NS-IBTS survey for the years 1991, 2001, 2011, and 2021.

### 3.2. Example 2: Derived survey variables

The following example demonstrates how `DATRASextra` can be used to derive standardised length-structured haul-level survey variables from relational DATRAS survey tables. Using a subset of Northeast Atlantic wolffish (*Anarhichas lupus*) survey observations included with the package, the workflow illustrates the aggregation of abundance and biomass catch rates into custom length classes, and estimation of stratified survey indices.

~~~
## Load the dataset
data(“wolffish”, package = “DATRASextra”)
## Clean the dataset
wolffish <-clean_datras(wolffish)
## Add swept-area estimates
wolffish <-add_swept_area(wolffish)
## Create numbers at length spectrum wolffish <-add_numbers_at_length(wolffish)
## Create weight at length spectrum wolffish <-add_weight_at_length(wolffish)
## Add total numbers per haul for two length classes wolffish <-add_total_numbers_by_haul(wolffish, length_cuts = c(0, 55, Inf))
## Add total weight per haul for two length classes wolffish <-add_total_weight_by_haul(wolffish, length_cuts = c(0, 55, Inf))
## Calculate stratified mean index numbers per swept area index <-calc_stratified_index(wolffish)
## Calculate stratified mean index biomass per swept area index_bio <-calc_stratified_index(wolffish, value_var = “HaulWgt”)
## Calculate spatial indicators indicators <-calc_spatial_indicators(wolffish)
~~~

Fine-scale number-at-length and biomass-at-length observations were first derived from the relational DATRAS tables and subsequently aggregated into user-defined length classes. We split the data set into two groups –approximately corresponding to juveniles and adults – using the length at 50% maturity (*L*_50_ = 55 cm) for wolffish. The resulting indices show variable but decreasing stratified mean abundance and biomass for both juvenile and adult wolffish (Figure 3). While these design-based stratified mean indices provide a simple and transparent summary of survey data and are useful for rapid exploration of trends, they do not account for factors such as changes in spatial distribution, variation in catchability, or other covariates that may influence survey catches. Consequently, model-based abundance indices are generally preferred for stock assessments and other applications requiring more precise and robust estimates of population trends [e.g. 15, 16, 17].

**Figure 3:**
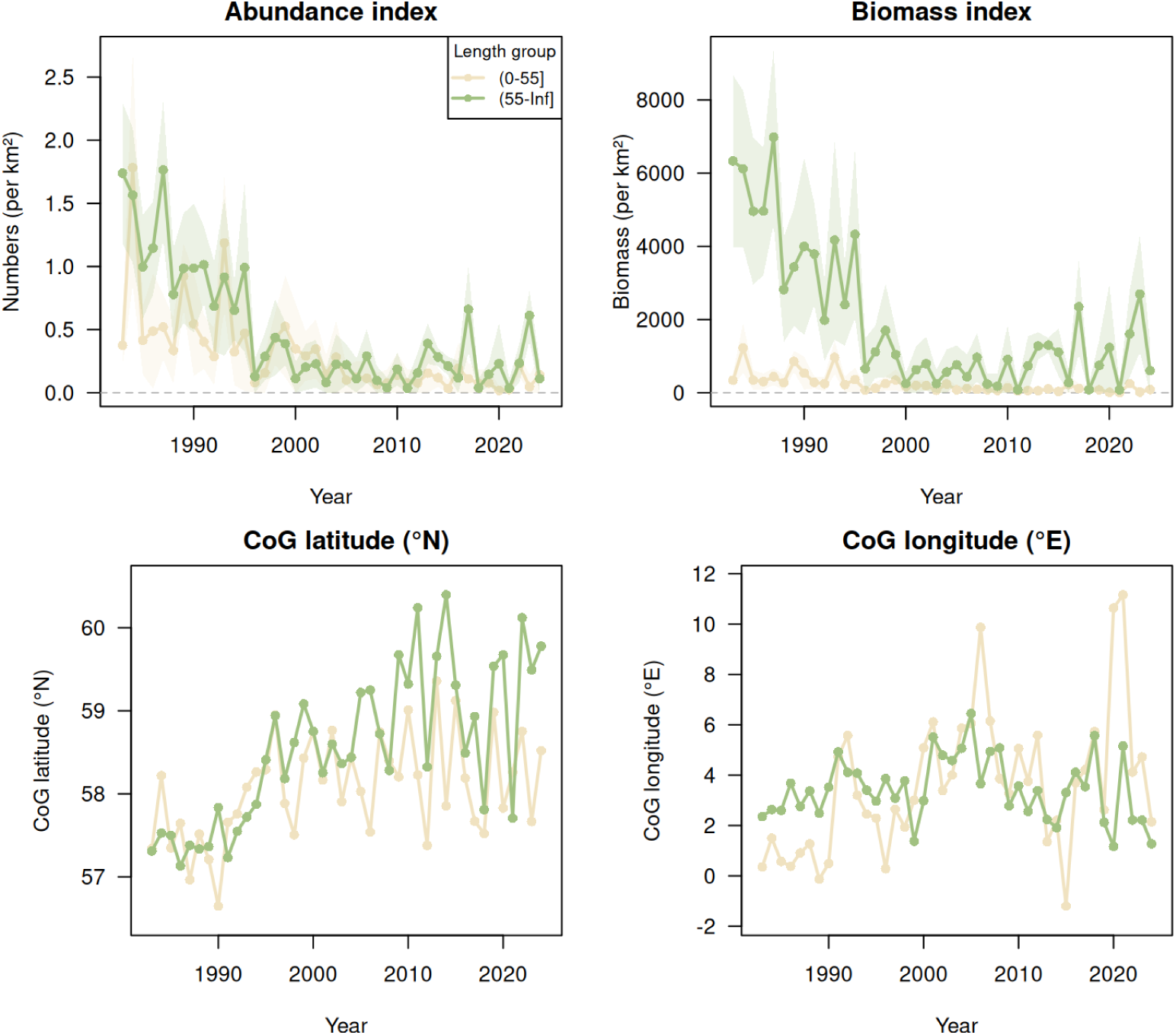
Example output generated with `DATRASextra` showing stratified mean survey indices for wolffish in the Northeast Atlantic in terms of abundance and biomass standardised by swept area and spatial indicators from 1983 to 2024. Shaded polygons represent 95% confidence intervals.

This workflow demonstrates how `DATRASextra` prepares standardised, length structured, haul-level data, indices, and indicators suitable for downstream stock assessment and ecological analyses that account for ontogenetic differences in distribution and abundance.

### 3.3. Example 3: Cross-survey integration and FishGlob workflows

The following example illustrates how `DATRASextra` can be used to reproduce key components of the FishGlob data-preparation workflow for integrated multi-survey analyses. Such workflows typically require substantial custom code to harmonise survey formats, derive standardised haul-level quantities, and generate analysis-ready datasets that can be combined across surveys and regions. For demonstration purposes, we use a small dataset with the same structure as the full workflow; however, the same steps can be applied directly to larger subsets or the complete DATRAS database. The example mirrors the procedures used to generate the Northeast Atlantic component of the FishGlob dataset [8].

~~~
## Load the dataset
data(“mini_fishglob”, package = “DATRASextra”)
dat0 <-mini_fishglob
## Clean the dataset
dat <-clean_fishglob(dat0)
## Reduce dataset size
dat <-prune_fishglob(dat)
## Compute swept area
dat <-add_swept_area_fishglob(dat)
## Estimate biomass
dat <-add_total_weight_by_haul_fishglob(dat)
## Format the FishGlob output fishglob <-as_fishglob(dat)
## Plot map with species richness plot_datras_overview(dat,
metric = “species_richness”,
by_year = TRUE,
multi_panels = TRUE)
~~~

The resulting datasets include haul-standardised quantities, survey metadata, and interoperable output formats for direct integration into downstream biodiversity and macroecological applications. Figure 4 shows an example visualisation of spatial and temporal variation in species richness (number of species) for the years 2015-2020. It was produced with the workflow shown above applied to a full snapshot of the DATRAS database (replacing `dat0 <-mini_fishglob` with `dat0 <-download_datras(years = 2015:2020)`), while omitting one survey each in Canada and the Mediterranean and the weight-by-haul step (which can drop species lacking length-weight parameters *a* and *b*). More details and a complete reproducible workflow for generating the Northeast Atlantic component of the FishGlob dataset are provided in the package FishGlob vignette and online documentation.

**Figure 4:**
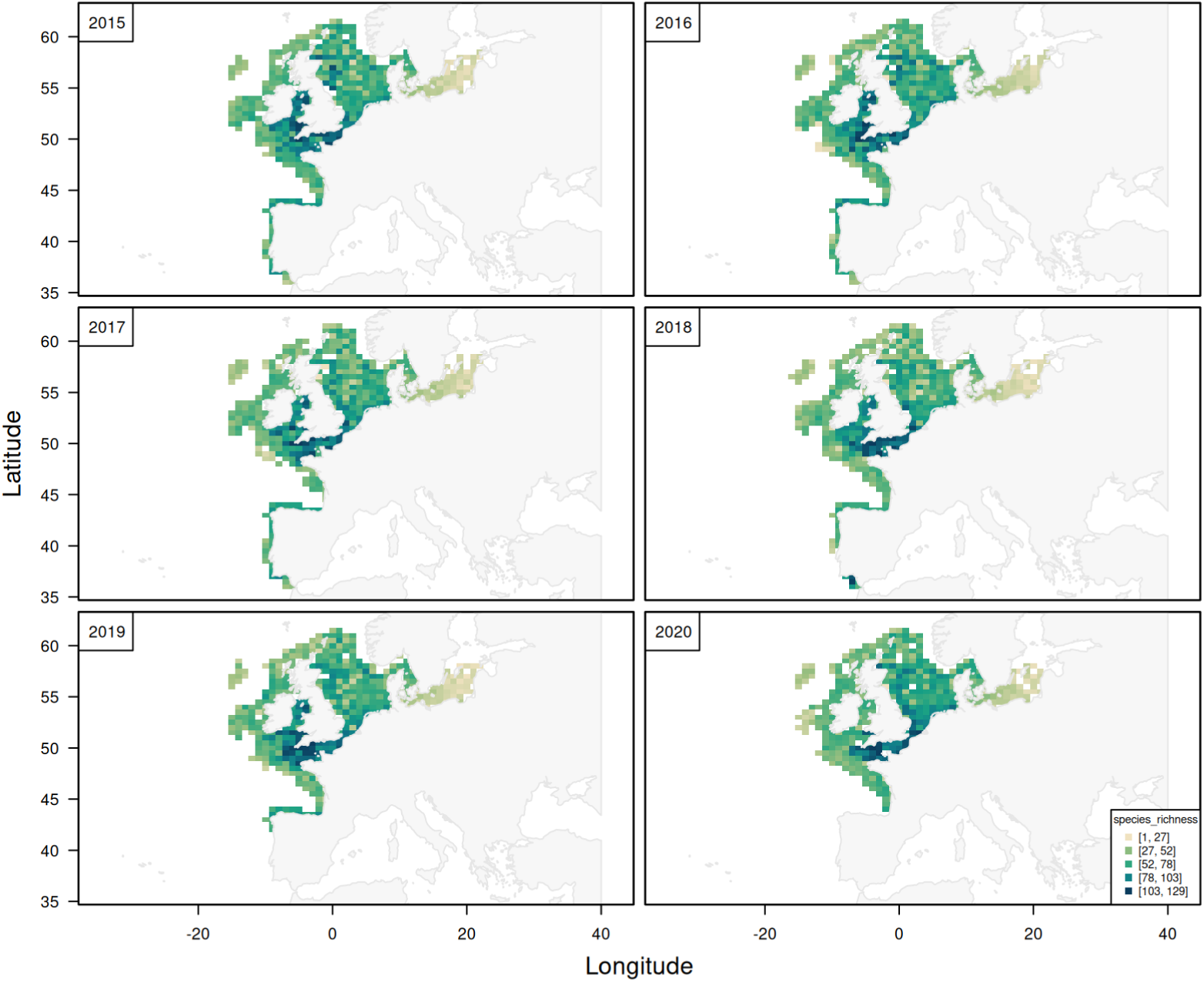
Example output generated with `DATRASextra` showing spatial and temporal variation in haul-level species richness (number of species) based on the ICES DATRAS database for 2015-2020. See the main text for the exact processing steps used to generate this figure.

## 4. Impact

`DATRASextra` improves accessibility, reproducibility, and standardisation of workflows based on ICES DATRAS survey data. By integrating data access, harmonisation, quality control, and generation of analysis-ready outputs into a single framework, the package substantially reduces the technical barriers for working with large and heterogeneous bottom-trawl survey datasets.

The package supports a broad range of scientific applications, including fisheries stock assessment, biodiversity monitoring, ecosystem analyses, species distribution modelling, life-history analyses, and large-scale macroecological syntheses. In particular, `DATRASextra` facilitates cross-survey integration and reproducible processing of long-term survey time series, enabling analyses that would otherwise require extensive project-specific scripting. For example, workflows used to reproduce the Northeast Atlantic component of the FishGlob dataset [8] can be implemented with only a few lines of code using `DATRASextra`, compared with thousands of lines of custom processing scripts previously required. The package is also being adopted within ICES expert groups: it is being used to investigate shifts in fish distribution and essential fish habitats (WKFISHDISH^1^) and the impact of warming on growth rates and fishery yields (WGGRAFY^2^).

By providing standardised and transparent workflows, the package also improves comparability across studies and institutions. This is particularly important for collaborative international research and advisory processes, where inconsistent preprocessing steps can otherwise lead to differences in derived survey products and scientific conclusions. The package incorporates established community recommendations and best-practice guidelines for bottom-trawl survey processing and cross-survey harmonisation [10], thereby supporting more consistent and reproducible analyses across users and applications.

`DATRASextra` has also simplified daily practice for users working with DATRAS data. Common but time-consuming tasks, such as downloading large survey subsets, merging relational tables, and preparing standardised outputs, are consolidated into a small number of interoperable functions. The package supports both interactive exploratory analyses and fully reproducible scripted workflows in R, making it suitable for research, advisory, and educational applications.

The software is released as an open-source R package under the GNU General Public License (GPL-3.0). Source code, issue tracking, and development history are publicly available on GitHub at https://github.com/tokami/DATRASextra. A dedicated pkgdown website [18] provides detailed documentation, tutorials, and end-to-end workflow examples at https://tokami.

github.io/DATRASextra. The package is actively developed and designed to accommodate future extensions, including improved downloading capabilities, additional visualisation tools, survey gear standardisation workflows, and expanded life-history and biological summaries derived from DATRAS data. All code and data used in this paper are provided to ensure full reproducibility.

## 5. Conclusions

DATRASextra consolidates the full DATRAS workflow, from raw data access to analysis-ready survey products, into a single reproducible R framework, extending rather than duplicating the capabilities of existing DATRAS-related software. As marine survey datasets continue to grow in size, complexity, and importance for ecosystem-based management, openly available tools that make their processing efficient, transparent, and reproducible become increasingly valuable. By lowering these barriers for one of the largest and most widely used bottom-trawl survey databases in the world, DATRASextra supports both routine survey processing and large-scale integrative analyses across fisheries and ecological research.

## Acknowledgements

We acknowledge ICES and the national institutes contributing survey data to DATRAS for maintaining and curating this essential resource [1]. We thank the developers of the icesDatras package and participants in ICES workshops and working groups (including WKFISHDISH2) for developing and sharing best-practice guidance for survey data processing and integration [11, 10]. We also acknowledge the developers and maintainers of the FishGlob dataset [8].

## 7. Author contributions

TKM: Conceptualisation, Funding acquisition, Investigation, Methodology, Project administration, Software, Validation, Visualization, Writing - original draft, Writing - review and editing; FM: Conceptualisation, Investigation, Methodology, Software, Validation, Writing - original draft, Writing - review and editing; CWB: Conceptualisation, Investigation, Methodology, Software, Validation, Writing - original draft, Writing - review and editing;

## 8. AI usage disclosure

OpenAI’s ChatGPT (GPT-5.2 Thinking; accessed March 2026) was used for copy-editing portions of the manuscript and suggesting wording and structure for roxygen documentation describing the package functionality. Anthropic’s Claude (claude-sonnet-4-6; accessed 2026) was used to assist with selected software development tasks, including bug fixes, refactoring, and code review. All AI-assisted outputs were reviewed, edited, and validated by the authors, who retain full responsibility for the accuracy, originality, licensing, and ethical and legal compliance of the manuscript, documentation, and software. Core software design, architecture, and the majority of implementation and testing were conducted by the authors independently of AI tools.

## Footnotes

1 https://www.ices.dk/community/groups/Pages/Wkfishdish3.aspx

2 https://www.ices.dk/community/groups/Pages/Wggrafy.aspx

